# Maternal omega-3 fatty acid deficiency affects fetal thermogenic development and postnatal musculoskeletal growth in mice

**DOI:** 10.1101/2022.10.13.512191

**Authors:** Vilasagaram Srinivas, Archana Molangiri, Saikanth Varma, Aswani Mallepogu, Suryam Reddy Kona, Ahamed Ibrahim, Asim K Duttaroy, Sanjay Basak

**Affiliations:** National Institute of Nutrition, Indian Council of Medical Research, Hyderabad, India; Department of Nutrition, Institute of Basic Medical Sciences, Faculty of Medicine, University of Oslo, Norway

**Keywords:** N-3 polyunsaturated fatty acid deficiency, Thermogenesis, Insulin growth factor, Uncoupling protein, Metabolism

## Abstract

Maternal omega-3 (n-3) polyunsaturated fatty acids (PUFAs) deficiency can affect offspring’s adiposity and metabolism by modulating lipid and glucose metabolism. However, the impact of n-3 PUFA deficiency on the development of fetal thermogenesis and its consequences is not reported. Using an n-3 PUFA deficient mice, we assessed fetal interscapular brown adipose tissue (iBAT), body fat composition, insulin growth factor-1 (IGF-1), glucose transporters (GLUTs), and expression of lipid storage & metabolic proteins in the offspring. The n-3 PUFA deficiency did not change the pups’ calorie intake, organ weight, and body weight. However, the offspring’s skeletal growth was altered due to excess fat to lean mass, reduced tibia & femur elongation, dysregulated IGF-1 in the mother and pups (p<0.05). Localization of uncoupling protein 1 (UCP1) in iBAT exhibited a reduced expression in the deficient fetus. Further, UCP1, GLUT1, *GPR120* were downregulated while FABP3, ADRP, GLUT4 expressions were upregulated in the BAT of the deficient offspring (p<0.05). The deficiency decreased endogenous conversion of the n-3 LCPUFAs from their precursors and upregulated *SCD1, FASN*, and *MFSD2A* mRNAs in the liver (p<0.05). An altered musculoskeletal growth in the offspring is associated with impaired browning of the fetal adipose, dysregulated thermogenesis, growth hormone, and expression of glucose and fatty acid metabolic mediators due to maternal n-3 PUFA deficiency. BAT had higher metabolic sensitivity compared to WAT in n-3 PUFA deficiency. Maternal n-3 PUFA intake may prevent excess adiposity by modulating fetal development of thermogenesis and skeletal growth dynamics in the mice offspring.

**Highlight:**

- Maternal n-3 PUFA deficiency dysregulated the development of fetal adipose browning
- N-3 PUFA regulates fetal thermogenic development by altering UCP1 expression
- BAT had higher metabolic sensitivity compared to WAT in n-3 PUFA deficiency
- Increased fat mass and IGF-1 played a role in promoting adiposity in n-3 PUFA deficiency

## 1. Introduction

Impaired thermogenesis arises from fetal adipose tissue dysfunction may impact early catch-up growth, offspring body composition [1], and predispose metabolic diseases in adult life. Maternal polyunsaturated fatty acids (PUFAs) during pregnancy correlate with the offspring’s body composition [2]. Postnatal n-3 long chain polyunsaturated fatty acids (LCPUFA) intake showed male-specific insulin-like growth factor-1 (IGF-1) changes in the offspring [3]. IGF-1 is a primary regulator for growth and body composition in fetal life and childhood, has a significant effect on the growth of the fetus and the placenta, as it is involved in the metabolism of muscle and fat tissue [4]. The concentration of IGF-1 in the mother’s blood correlates with the newborn’s Ponderal index [5]. However, the potential implications of n-3 fatty acid deficiency during pregnancy on IGF-1 concentration and its relationship in mother and offspring are unknown. The maternal, fetal, and neonatal LCPUFAs status can determine the metabolic outcome in infancy and later life.

Unlike essential nutrients, LCPUFAs are straightforward to consume, but the right balance of n-6/n-3 fatty acids is important for proper growth, adiposity, and insulin sensitivity [6]. Differential effects of maternal n-6 to n-3 fatty acids on offspring’s adiposity and its relation with obesity are suspected [7]. The optimal n-6 to n-3 fatty acid regulates adiposity in diverse ways, including a change in thermogenesis [8-13], skeletal integrity and bone development [14-17], glucose metabolism [18-20], hormonal changes [21]. The n-3 PUFAs regulate lipid storage and metabolism by decreasing the esterification of the lipids [22]. These fatty acids promote lipids oxidation and decrease triglyceride synthesis by increasing fatty acids oxidation, decreasing tissue lipid storage in the liver. The higher n-6 fatty acids produce arachidonic acid (ARA)-derived metabolites that promote adipogenesis [23], inhibits white adipose tissue (WAT) to brown adipose tissue (BAT) conversion [24].

Thermogenesis control energy allocation by modulating oxidation and storage of fats. Browning white adipose depots and activation of uncoupled respiration in brown fat lowers the fat accumulation. Brown fat is indispensable for maintaining the neonate’s body temperature, and it helps in the burning of white fats for meeting the energy of the growing neonates. WAT sparsely intermingled with brown fat-like cells known as brite or beige adipocytes [25, 26]. Uncoupling protein-1 (UCP1) is a prominent thermogenic protein involved in browning. The beige fat regulates glucose metabolism independent of UCP1, thus controlling systemic energy metabolism and glucose homeostasis [27]. Browning agents regulate energy homeostasis concerning thermogenesis. Enhancing brown adipogenesis prevents metabolic dysfunction in the offspring mice [28]. The functional ability of n-3 PUFA in activating brown fat thermogenesis is promising [11]. N-3 PUFA-induced thermogenesis is modulated by multiple mechanisms including gut-mediated energy expenditure [29], brown-specific microRNA (miRNA) upregulation [30], upregulation of thermogenesis biomarkers. Biomarkers of thermogenesis, such as protein related domain containing 16 (PRDM16), peroxisome proliferator-activated receptor-gamma coactivator 1 alpha (PGC1α) and UCP1 increased considerably in BAT from eicosapentaenoic acid (EPA)-fed mice compared to HFD.

In contrast, these markers were not detectable levels in either the subcutaneous or visceral WATs [13]. The thermogenic response to β3-adrenergic stimulation was enhanced in BAT and WAT by a diet rich in low n-6 to n-3 fatty acids [8]. N-3 PUFAs stimulate the beige and brown adipocyte differentiation by activating G-protein coupled receptor 120 (GPR120), which induces the growth hormone fibroblast growth factor 21 [31]. By regulating differentiation, n-3 PUFA can remodel fetal adiposity by balancing fat storage, and its oxidation [32]. Despite these data, the effects of n-3 PUFA deficiency in the browning of fetal adipose and its postnatal effects in adiposity are not studied.

Inadequate n-3 PUFA leads to docosahexaenoic acid (DHA) deficiency that could affect fetal and postnatal neurodevelopment [33], feto-placental changes in epigenetics [34], offspring growth and lipogenic capacity [35], and others. PUFA and its metabolites also regulate bone formation and resorption [17]. Diet rich in n-3 PUFA is reported to prevent bone loss [15]. However, limited data is available on maternal n-3 PUFA deficiency and its earliest changes on the skeletal growth in the offspring. India has the second-largest number of obese children globally [36], where fetuses are often nurtured in n-3 fatty acid deficiency due to its lower intake during pregnancy and lactation [37]. Maternal n-3 PUFA depletion during *in utero* development of adipose browning, its postnatal effects on musculoskeletal development, and IGF-1 of the offspring are not known yet. Moreover, the mechanism of n-3 PUFAs on the induction of browning response or increase in the activity or amount of UCP1 in developing adipose tissue is not clear.

We hypothesized that altered skeletal growth could be due to body fat changes and impaired thermogenic fat development during fetal adipogenesis. Using a recently developed n-3 PUFA deficient mouse model [34], we measured UCP1-mediated adipose browning in a fetus and the relationship of IGF-1 in mother-pups, expression of glucose, and lipid metabolic transporters in adipose and livers of offspring mice.

## 2. Materials and Method

### 2.1 Diets, animals and sampling

All the procedures involved in the animal experiment were conducted in accordance with the guidelines of the committee for the control and supervision of experiments on animals (CPCSEA), Government of India. The study was approved by the institutional animal ethical committee of the National Institute of Nutrition, Hyderabad, India (No. NCLAS/IAEC/02/2017). The fetus and offspring examined in the present study are the continuity of our recently published report [34].

Weanling female (n=30 per group) Swiss albino mice were caged in pairs at 23±3°C, 55% ± 10% relative humidity with ∼12h light/dark cycle and fed on n-3 PUFA sufficient and deficient diet ad libitum for five weeks before the introduction of an age-matched mating partner. Experimental diets were formulated as per the AIN93 diet with modification of fat composition. The feed’s dietary and fatty acid composition are similar in the present study, as mentioned in our previously published work [34]. Diet consists 90% of base mixture [carbohydrate (54.5%), protein (25%), cellulose (5%), salt mixture (4%), vitamin mixture (1%), L-cystine (0.3%) and choline chloride (0.2%)] and 10% of fat source. Peanut and palmolein oils were used to reflect the Indian dietary fat intake since these are widely consumed oil sources, while linseed oil was used for the n-3 PUFA source. Two different blends of peanut oil, palmolein oil, and linseed oil were used as fat sources and mixed with base mixtures to obtain the n-6: n-3 PUFA ratios (50: 1 and 2:1) in the experimental diets. The isocaloric diets were formulated to ensure that each mouse received the same calories and nutrients but different proportions of n-3 and n-6 fatty acids in the total mixture of PUFA. Blended oils were flashed from time to time with nitrogen gas to prevent oxidation.

A line diagram of dietary intake and data collection time points of the study is presented in **Supplementary Fig.1**. Pregnant (gD 14.5-17.5) mice (n=6) were euthanized by cervical dislocation, and uterine horns bearing implanted fetus were processed on ice to collect the fetus. Since BAT and bone development closely overlap with gD 14.5-17.5, a mouse fetus was investigated during this window [38]. Collected fetuses (n=6 fetus/each) were immediately stored in RNA later, formaldehyde (4%) solution and snap-frozen to carry out mRNA expression, immunohistology/fluorescence, and protein expression, respectively. The remaining pregnant mice were allowed until parturition. Blood was collected from the retro-orbital plexus of the offspring and was euthanized to collect liver and fat (white and brown) tissues. The litter size of pups was normalized for each group. The pups’ food intake and body weight were monitored throughout the study. Food intake of the offspring was calculated daily by subtracting leftover diet from the initial supplied and expressed in grams/week of each group. Feed efficiency (%) was calculated as (mean body weight gain *100)/energy intake, whereas energy intake (kcal/day) = mean food consumption x dietary metabolizable energy (kcal).

### 2.2 Fatty acid profiles

Total lipids were extracted from plasma (100µl) as described before [39]. Samples were subjected to methylation at 70°C with 2% methanolic sulphuric acid (containing butylated hydroxytoluene 10 mg/L) for 4 hours. Methyl esters of fatty acids (FAME) were separated out of the fraction after cooling. Gas chromatography (Perkin Elmer Clarus 680) was performed using a flame ionization detector and a fused silica capillary column (#24019, Supelco, Bellefonte, PA, USA) to determine fatty acid composition of FAME fraction. To quantify total fatty acid content, C17:0 fatty acid was used as an internal standard and added to the samples before methylation. Individual fatty acid peaks were identified by comparison with the 63B standard (Nu-chek-Prep, USA).

### 2.3 Assessment of musculoskeletal development and lower limb analysis by dual-energy X-ray absorptiometry (DEXA)

DEXA (Discovery, Hologic, Bedford, MA, USA) was used to evaluate the body composition i.e., bone mineral density (BMD), bone mineral content (BMC), body mass, and whole-body fat distribution of all mice. Prior to the DEXA scan, the mice were weighed and injected intraperitoneally using ketamine and xylazine at a final dosage of 80 and 10 mg/kg of body weight, respectively. The mice were lying flat on the scanning bed, with their limbs and tails protruding from their bodies. Total fat, fat percentage, lean body mass, bone mineral density (g/cm2), mineral content (g), were obtained. The lengths of the tibia and femur were quantified in pixels using DEXA radiographs using ImageJ 1.50i (NIH, USA).

### 2.4 Plasma insulin growth factor analyses

Plasma IGF-1 level was analyzed using ELISA assay (#E-EL-M3006, Elabscience, USA). Briefly, 100μl of standard and diluted samples were incubated in a pre-coated 96-well plate for 90 min. The samples were coated with a biotinylated detection antibody for 1h followed by incubation with HRP conjugate for 30min, and substrate reagent for 15min. The enzyme action was terminated by a stop solution (50μL) and OD was measured at 450nm. A four-parameter logistic curve was constructed to analyze the concentration of plasma IGF-1 as per the supplier’s guideline.

### 2.5 Immunofluorescence of UCP1in fetal interscapular brown adipose tissue

Fetuses were collected from 14.5-17.5gD pregnant mice and fixed in 4% paraformaldehyde. After fixing, tissues were dehydrated and embedded in paraffin. Then sagittal sectioning of interscapular tissue was performed to obtain 4µm sections and coated in slides. The sections were de-paraffinized by placing the slides on a hot plate for 1h at 60°C. De-paraffinized slides were immediately washed in xylene to prevent any further solidification. The slides were rehydrated stepwise with 95%, 75%, and 50% alcohol about 10 min each and finally washed with distilled water. The antigen retrieval step was performed by transferring the slides to 10mM citrate buffer (pH 6.0) in the microwave oven for 5 min. After two washes with 20mM PBS, the slides were blocked with 3% horse serum and kept in a humidified chamber for 90 min. Next, the slides were washed gently two times with 20mM PBS for 10 min and incubated overnight at 4°C with primary antibody (anti-UCP1, CST #14670s; 1:50 dilution), followed by incubation with Alexa fluor goat anti-rabbit IgG (#A11034, Invitrogen;1:200 dilution) for 1h in a humidified chamber. Slides were washed with 20 mM PBS twice after incubation and mounted with DAPI solution (#F6057, Sigma). The images were captured under 40X magnification using a fluorescent microscope (Leica Microsystem, Germany) and quantified for total fluorescence with Image J software. Corrected total cell fluorescence (CTCF) was calculated by formula; CTCF=Integrated density – (area of selected cell x mean fluorescence of background readings) after measuring fluorescence by ImageJ 1.50i (NIH, USA) and expressed in arbitrary units.

### 2.6 Measurement of protein expression by immunoblotting

Adipose tissues (white and brown) were finely ground using a mortar and pestle (in liquid nitrogen) followed by the addition of 500 µl of radioimmunoprecipitation assay (RIPA) buffer (protease inhibitors included). Lysates were spun at 12000g for 20min at 4°C following sonication at 30% amplitude for 10s twice on ice. A total of 50-100 µg of proteins/lanes was used in immunoblotting after protein estimation by BCA assay (Pierce, USA). Proteins were resolved by SDS-PAGE (10%) before their transfer to polyvinylidene difluoride membranes (Immobilon-P, Millipore Corp.). After blocking (#1706404, Bio-Rad), membranes were immunoblotted with anti-UCP1 (1:1000, CST 14670s), anti-GLUT1 (1:1000, CST 12939s), anti-GLUT4 (1:2000, Thermo Fisher-PA5-23052), anti-ADRP (1:1000, Thermo Fisher-PA1-16971), anti-FABP4 (1:1000, Thermo Fisher-PA5-3059), anti-FABP3 (1:4000, Adipogen-AG25A0040) and anti-β-actin (1:1000; ab-8227, Abcam) antibodies. Blots were incubated for 12h in a cold room followed by 1h incubation with HRP conjugated anti-rabbit/anti-mouse IgG antibody. The blots were detected using ECL substrate (#1705061, Bio-Rad), signals were recorded by G-box chemiluminescence imaging system (Syngene, USA) and quantified by ImageJ 1.50i (NIH, USA).

### 2.7 Expression of genes by RT-qPCR

Tissues (adipose, liver) were finely minced (50mg) and transferred into 1 ml of cold Tri reagent (# T9424 Sigma) containing 2 mm zirconia beads (#11079124Zzx, Biospec). Tissues were homogenized using bead beater for 1-2 min (Mini Bead beater, Biospec) and spun at 12,000 g for 10 min to eliminate cell remnants (4°C). Total RNA was purified as per manufacturer instructions. Isolated total RNA was undergone gDNA elimination (#AMPD1, Sigma) followed by quality (260/280 and 260/230 ηm) and quantity checked by spectrophotometer (#ND1000, Thermo Scientific). Total RNA (1µg) was converted to cDNA as per the manufacturer’s instruction (#1708891 Bio-Rad). Relative mRNA expression was performed using cDNAs as a template with pre-designed primers compatible with SYBR green I chemistry (gene ID, pathways, and nucleotide sequences are enclosed as **Supplementary Table 1**). The mRNA expression was performed in FAST mode using a real-time PCR machine (Quant5, Life Technology, USA). The reactions setup was carried out in a microamp optical 384-well plate (Life tech. #4309849384) using the power SYBR green PCR master mix (Life tech. #4367659). After a run, the Ct (threshold cycle) values were used to calculate the mRNA expression by using the comparative delta Ct method (2^−ΔΔCt^). Data were presented as fold change over control after normalizing with the endogenous reference gene.

**Table 1.** Plasma total fatty acid composition (% nmole) of the 21-d pups born to dam fed with n-3 deficient (0.13en%) and n-3 sufficient (2.26en%) diet

| Fatty acids<br>(nmol %) | n-3 sufficient<br>(n=5) |  | n-3 deficient<br>(n=5) |  | p value | Fold<br>change<sup>#</sup> |
| --- | --- | --- | --- | --- | --- | --- |
|  | Mean | SEM | Mean | SEM |  |  |
| C16:0<br>(Palmitic acid) | 27.49 | 1.0 | 25.70 | 1.09 | 0.259 | - |
| C18:0<br>(Stearic acid) | 19.28 | 1.20 | 21.72 | 0.72 | 0.135 | - |
| C18:1<br>(Oleic acid) | 11.67 | 0.63 | 11.59 | 0.72 | 0.939 | - |
| C18:2 n-6<br>(Linoleic acid) | 15.63 | 1.32 | 15.34 | 0.46 | 0.842 | - |
| C20:4 n-6<br>(Arachidonic acid) | 8.23 | 0.49 | 16.63 | 0.94 | **<0.0001\*\*\*** | 2.0 (+) |
| C22:4 n-6<br>(Docosatetraenoic acid) | 4.21 | 0.47 | 3.70 | 0.36 | 0.435 | - |
| C18:3 n-3<br>(α-Linolenic acid) | 0.96 | 0.10 | ND | ND | - | - |
| C20:5 n-3<br>(Eicosapentaenoic acid) | 0.53 | 0.24 | ND | ND | - | - |
| C22:6 n-3<br>(Docosahexaenoic acid) | 8.56 | 0.76 | 1.88 | 0.10 | **<0.0001\*\*\*** | 4.5 (-) |
| <sup>1</sup>Σn-6 PUFA | 12.44 | 0.96 | 20.33 | 1.3 | 0.08 | 1.6 (+) |
| <sup>2</sup>Σn-3 PUFA | 9.09 | 1.0 | 1.88 | 0.10 | **0.0129\*** | 4.8 (-) |
| ΣLC(n-6): (n-3) | 1.3:1 | - | 10.8:1 | - | - | 7.9 (+) |

<sup>#</sup> Fold/folds change in fatty acid concentration is depicted by increase (+) or decrease (-) over n-3 sufficient group. ND = non-detectable.
<sup>1</sup> $\Sigma$ LC n-6 PUFA is the sum of C20:4 n-6 and C22:4 n-6
<sup>2</sup> $\Sigma$ LC n-3 PUFA is the sum of C20:5 n-3 and C22:6 n-3
Student's t-test between n-3 deficient and n-3 sufficient group; Mean ± SEM, \* p<0.05; \*\*\* p< 0.0005;

### 2.8 Statistical analyses

The student’s t-test was performed to compare two groups using Graphpad Prism v.8. Statistical significance was considered when the p-value was less than 0.05. The experiments were performed independently with repeated times, as indicated in the text or figure legends. The experimental values are expressed as a mean ± standard error of the mean (SEM).

## 3. Results

### 3.1 Food intake, feed efficiency, and bodyweight of the offspring

The cumulative food intake (g) of the n-3 PUFA deficient and sufficient fed offspring did not differ significantly (week 1: n-3 def. vs n-3 suff. = 39.64 ± 5.41 vs. 40.12 ± 3.95 to week 9: 475.78±17.85 vs. 466.22 ±28.58, p>0.05) over a period of nine-weeks. No significant changes were observed in feed efficiency (%) between these groups (n-3 sufficient vs. n-3 deficient: 1.11 ± 0.22 vs. 1.17 ± 0.27, p>0.05). Both groups were comparable in overall and sex-specific body weight in the pups (p>0.05, **Supplementary Table 2**). There was no difference in the liver or adipose weights between these groups (data not presented).

### 3.2 Plasma fatty acid composition in mice offspring

Gas chromatography was used to analyze the fatty acids composition in the plasma of 21 d old mice (**Table 1**). The arachidonic acid,20:4n-6 (ARA) levels were considerably increased by ∼2 folds in n-3 deficient pups (p<0.05), but linoleic acid, 18:2n-6 (LA) levels remained unchanged in both, indicating increased conversion of LA to ARA in n-3 PUFA deficient mice compared to its counterpart. The presence of docosahexaenoic acid,22:6 n-3 (DHA) was significantly reduced by ∼4.5 folds (p<0.05) in n-3 PUFA deficient mice’ plasma. The total n-3 PUFA content was substantially improved by ∼4.8 folds in n-3 sufficient offspring (p<0.05). The overall proportion of n-6 to n-3 PUFAs in the pup’s plasma from both groups was closely similar with placental n-6 to n-3 PUFAs ratio [34], indicating that dietary deficiency of n-3 PUFA was linearly transferred from mother to the pups, as evidenced by the plasma fatty acids composition.

### 3.3 Musculoskeletal growth and adiposity of the offspring

N-3 PUFA deficient pups showed a substantial increase in the fat mass and fat percentage compared to pups fed with n-3 PUFA sufficient diet (n-3 suff. vs. n-3 def.: fat mass: 5.033 ± 0.57 vs. 6.810 ± 0.52, p=0.034, and fat percentage: 20.29 ± 1.56 vs. 24.65 ± 1.158, p=0.036, **Fig. 1a**). The total body mass showed an upward trend for n-3 deficient pups but the changes were insignificant between groups (n-3 suff. vs. n-3 def.: body mass: 24.59 ± 1.42 vs. 27.47 ± 1.389, p=0.166, **Fig. 1a**). Repeated DEXA measure of male 21d offspring (n=6) showed a significant morphological difference in limb elongation (**Fig.1b)**, although there were no significant changes in bone mineral density (BMD) and bone mineral content (BMC) in these mice (data not shown). Region of interest was used to measure the length of femur and tibia after calibrating DEXA radiograph with Image J software and expressed in pixels. The total leg, femur, and tibia length were significantly decreased in n-3 deficient pups (n-3 suff. vs. n-3 def.: tibia (pixels): 0.708 ± 0.012 vs. 0.618 ± 0.04, p=0.034; femur (pixels): 0.59 ± 0.02 vs. 0.49 ± 0.03, p=0.014; leg total (pixels): 1.298 ± 0.03 vs. 1.106 ± 0.07, p=0.019, **Fig. 1c**). Since IGF-1 hormonal level is a growth indicator, its expression and concentration were evaluated to follow the growth trajectory during n-3 fatty acid deficiency. The mRNA expression of the *IGF-1* was significantly increased by ∼1.7 folds (**Fig.1d**, p=0.003) in the n-3 deficient WAT of the 21d pups. N-3 PUFA deficiency significantly lowered the plasma IGF-1 concentration in dams during pregnancy [n-3 suff. vs. n-3 def. (ng/ml): 1819 ± 110.9 vs. 1507 ± 58.69, p<0.05, **Fig. 1e**]. In contrast, IGF-1 concentrations were significantly increased in n-3 deficient pups [n-3 suff. vs. n-3 def. (ng/ml): 2493 ± 77.87 vs. 2691 ± 45.53, p<0.05, **Fig.1e**]. All these data indicate n-3 PUFA deficiency induces a disruption in the pup’s body fat distribution, musculoskeletal growth, IGF-1 levels in dams and pups.

**Fig. 1.**
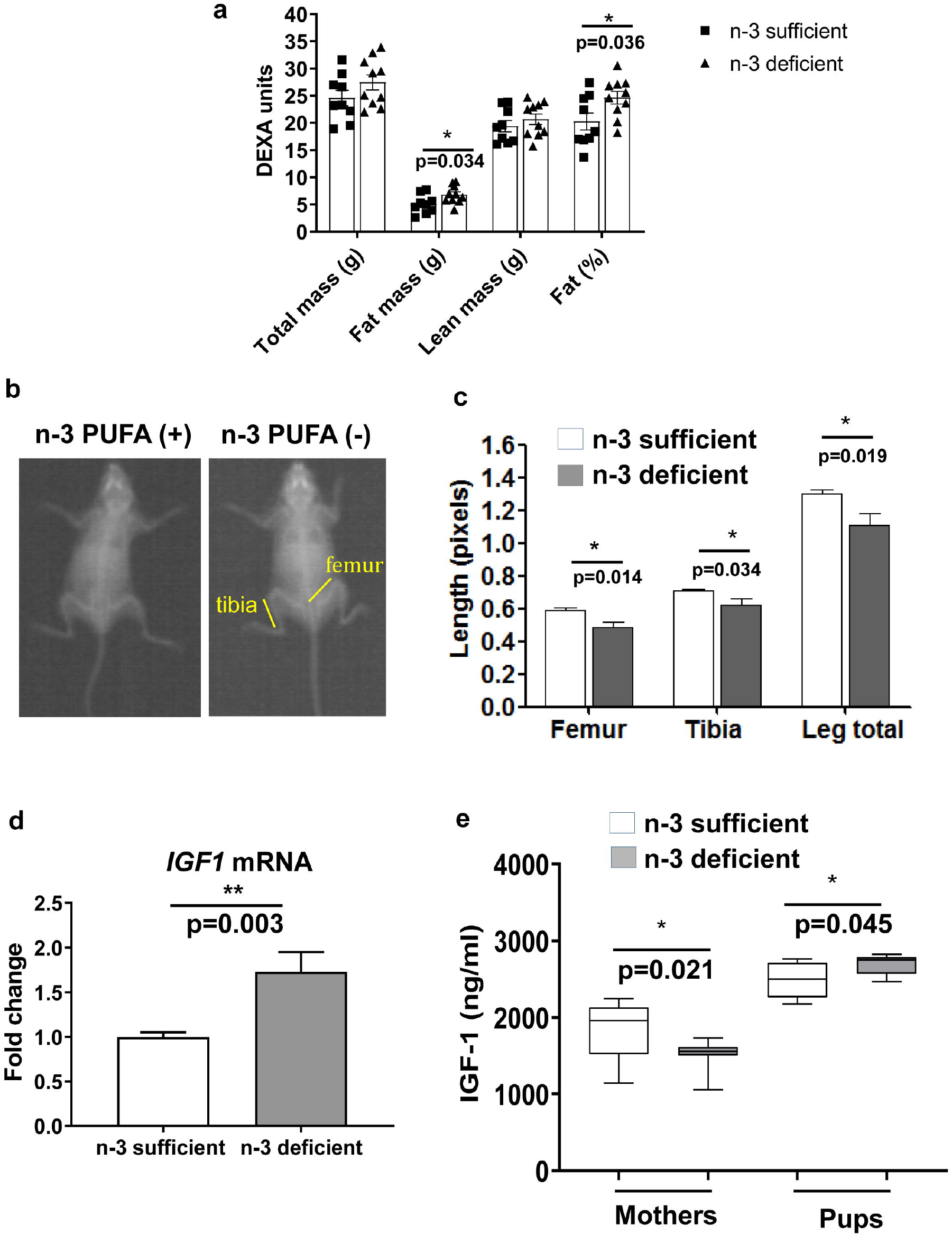
Fat and lean mass distribution, lower limb analysis, and insulin growth factor-1 in 21d offspring, whose mothers were fed on n-3 PUFA sufficient and deficient diet. **a**. Body mass and fat mass distribution (n=9-10) was measured by DEXA (dual-energy X-ray absorptiometry). Mean of total mass, fat mass and lean mass are expressed in grams, while the fat content is expressed in percentage. **b**. A representative DEXA image and anatomy of the lower limb of 21d male pups. **c**. DEXA radiographs analysis of lower limbs, expressed in pixels (n=6). **d**. Expression of IGF-1 mRNA (n=6) in white adipose tissue presented as fold change of expression over n-3 PUFA sufficient group after normalized with endogenous control. **e**. Plasma IGF-1 of mothers (n=12) and pups (n=8). Values are expressed in mean ± SEM. * p < 0.05 is considered as significant vs. n-3 sufficient (Student’s t-test).

### 3.4 Development of fetal brown adipose tissue in n-3 PUFA deficiency by scoring expression, and localization of uncoupling protein-1

To determine whether altered musculoskeletal growth and fat mass in n-3 deficient pups were due to impairment of brown adipogenesis during fetal development, we evaluated the expression and localization of a predominant brown adipose tissue marker, uncoupling protein 1 (UCP1) in tissues collected from the fetus’s interscapular region (**Fig.2a**). Normalized UCP1 expression was calculated in terms of corrected total cell fluorescence (CTCF). The CTCF of UCP1 was significantly reduced by ∼1.5 folds in the fetus of n-3 PUFA deficient mice (n-3 PUFA suff. vs. n-3 PUFA def.: 0.90 ± 0.083 vs. 0.57 ± 0.096, p=0.013, **Fig.2 b-c**). Since the development and transition of fetal BAT are regulated through UCP1, its expression was measured longitudinally in the offspring BAT. The UCP1 expression was significantly decreased by ∼2.9 folds (p<0.0001) in n-3 PUFA deficient 21d-offspring (**Fig.3 a-b**). A significantly reduced expression of UCP1 in the fetus and offspring indicated an altered thermoregulation and energy metabolism during n-3 PUFA deficiency.

**Fig. 2.**
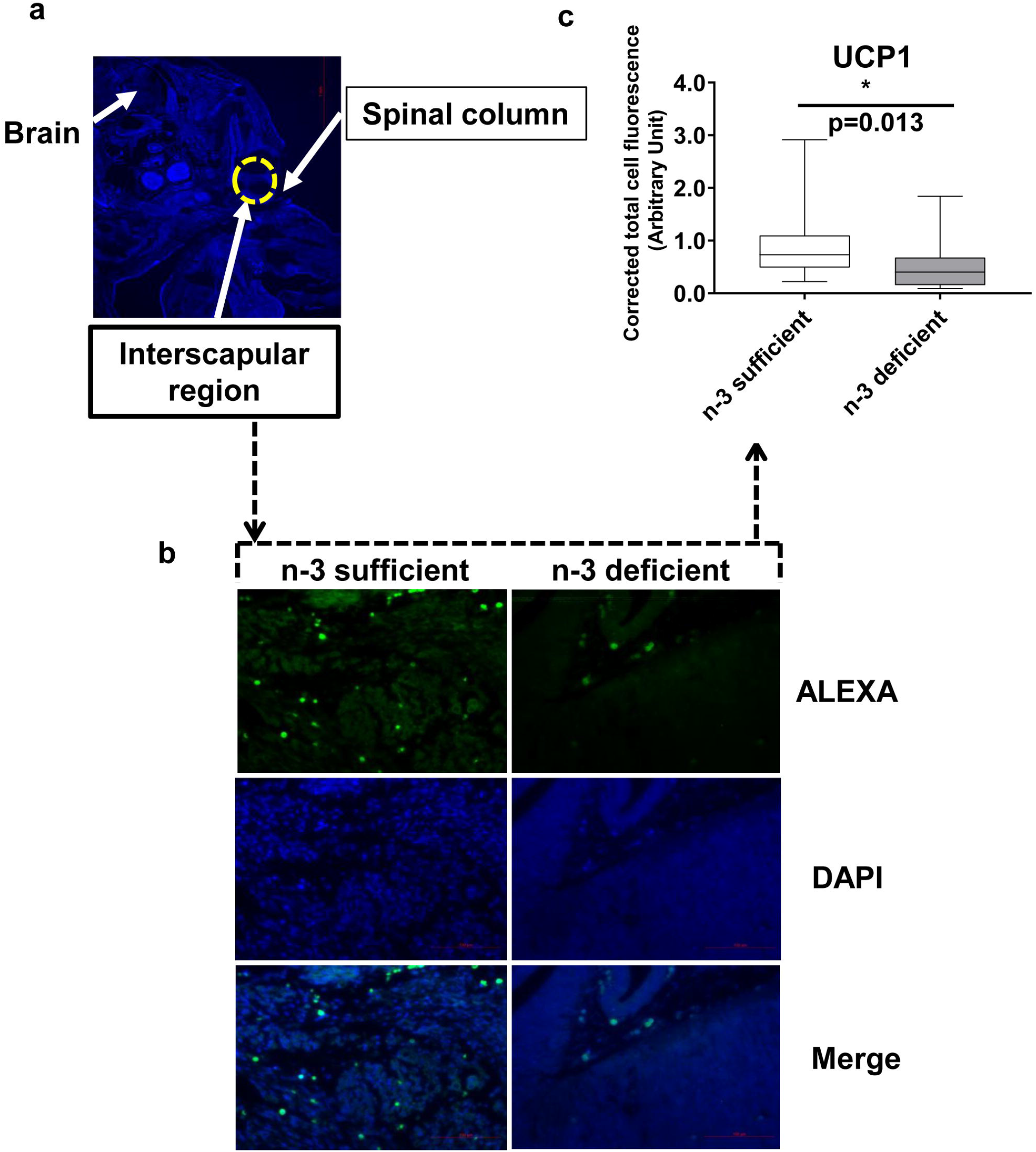
UCP1 localization and expression in foetal tissue by immunofluorescence. Foetal tissues were obtained from n-3 sufficient and deficient fed pregnant mice (14.5-17.5gD) (n=6/group). Tissues were fixed in 10% formalin, parrafinised, deparaffinised, and stained with rabbit anti UCP1 protein (primary) followed by Alexa flour-488 goat anti-rabbit antibody (secondary). Images were captured by LEICA laser capture microscope. **a**. Cross-sectional image of whole fetus showing brain, spinal column, interscapular region (magnification 10x). **b**. Immunofluorescence staining of UCP1 protein (green), nuclei (blue) of fetal interscapular tissue at 40x magnification. **c**. Corrected total cell fluorescence (CTCF) was calculated after measuring the fluorescence (CTCF= integrated density – (area of selected cell x mean fluorescence of background readings) and expressed in arbitrary units as mentioned in the method. Values are expressed in mean ± SEM. * p<0.05 vs. n-3 sufficient (Student’s t-test).

**Fig. 3.**
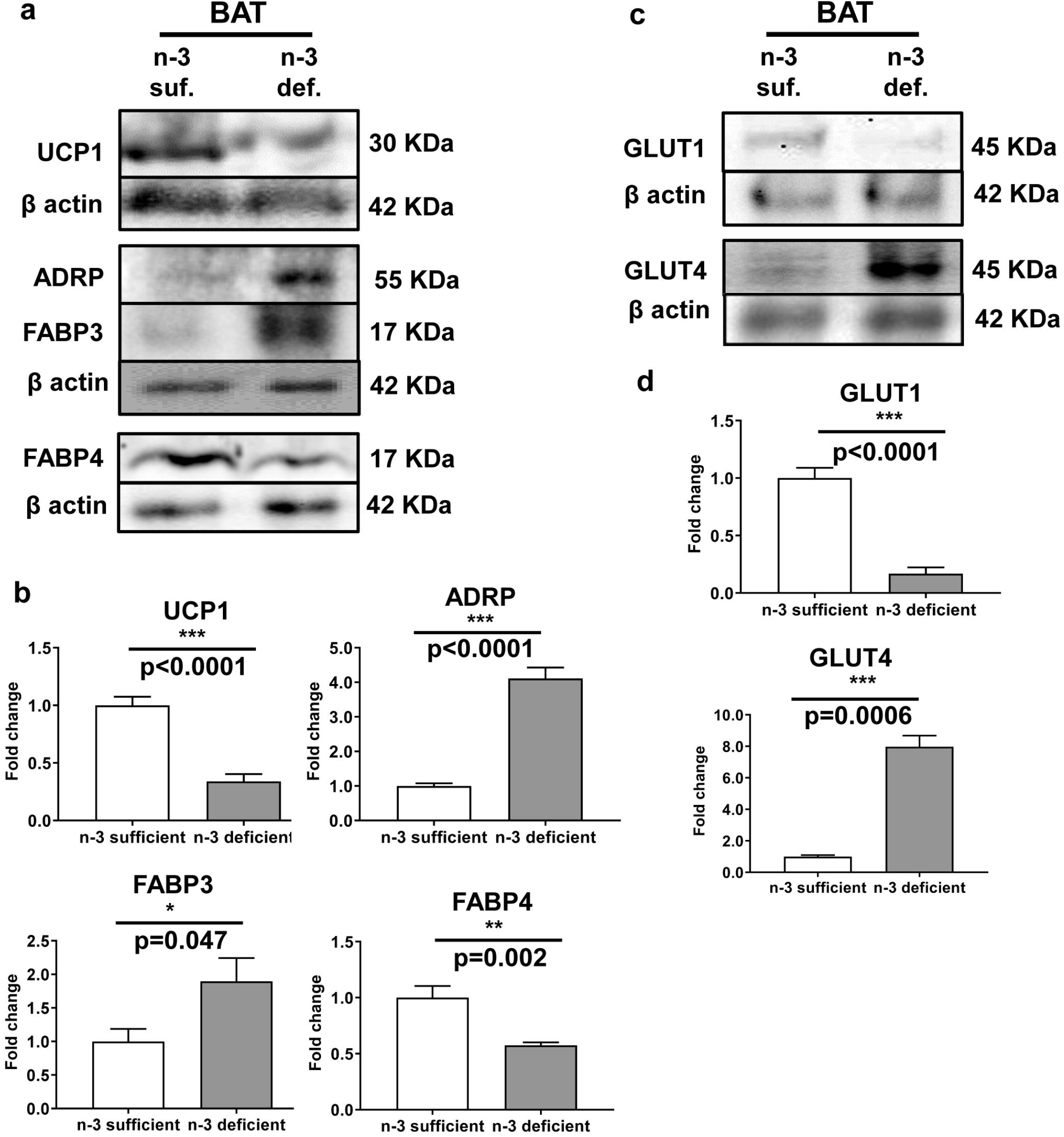
Effect of n-3 PUFA deficiency on the expression of (**a-b**) thermogenesis mediators, lipid storage & binding proteins, and (**c-d**) glucose transporters in brown adipose tissue (BAT) of 21d male pups born to n-3 PUFA sufficient and deficient dams. Expression levels were evaluated by immunoblotting of the proteins of the BAT homogenate obtained from mice (n=4 per group) and normalized the levels by the expression of β-actin. The expression level was reported as relative density from the three independent (n=12) experiments. Data are expressed in mean ± SEM. Individual p-values are indexed in figures. * p<0.05, ** p<0.005; *** p<0.0001 vs. n-3 sufficient diet (Student’s t-test).

### 3.5 Effect of maternal n-3 PUFA deficiency on the expression of energy metabolism mediators in pups’ brown adipose tissue

Since brown adipose stores metabolic energy as triacylglycerol in lipid droplets (LDs), expression of LD-associated proteins such as perilipin 2 or ADRP, FABP4, FABP3 were measured in offspring’s BAT. Expression of ADRP and FABP3 were significantly upregulated by ∼4.1 folds (p<0.0001) and ∼1.9 folds (p=0.047), while FABP4 expression was decreased by ∼1.7 folds (p=0.002) in n-3 deficient BAT (**Fig.3 a-b**). Since the expression of glucose transporters (GLUTs) is associated with thermogenesis, the predominant GLUT mediators such as GLUT1 and GLUT4 were analyzed in the BAT. The GLUT1 expression was decreased significantly by ∼5.9 folds (p<0.0001), whilst GLUT4 expression was increased by ∼7.9 folds (p=0.0006) in the n-3 deficient BAT **(Fig.3 c-d)**.

Dietary PUFAs and their metabolites may affect glucose and lipid metabolism in diverse ways including gene expression. Therefore, mRNA expression of tissue-specific functional genes (**Supplementary Table 1)** was measured in BAT. The mRNA expression of *UCP1* was downregulated by ∼1.6 folds (**Fig.4a**), *GPR120* by ∼4.6 folds (**Fig.4b**), *FGF 21* by ∼1.6 folds (**Fig.4c**), *GLUT1* by ∼4 folds (**Fig.4d**), and FABP3 (**Fig.4f**) were upregulated by ∼3.1 folds in BAT of n-3 deficient offspring while the expression of *PRDM16, PGC1*_α_, *GPR40, GLUT3, GLUT4, FABP4* did not change in both groups. The BAT mRNA expression of glucose transporter, fatty acid-binding protein, and free fatty acid receptor was significantly altered in n-3 PUFA deficiency.

**Fig. 4.**
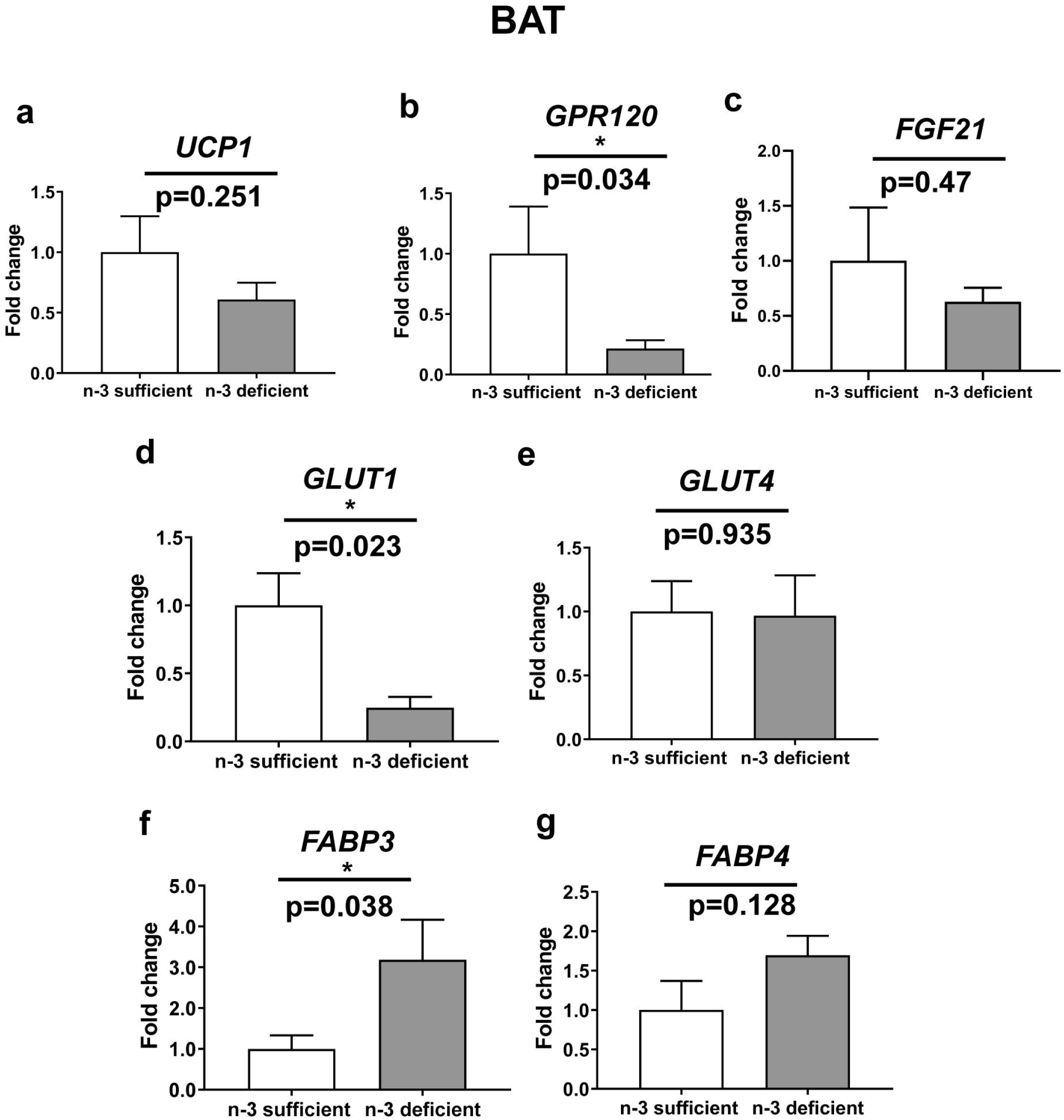
Effect of n-3 PUFA deficiency on the expression of genes involved in thermogenesis mediators, glucose transporters, and fatty acid-binding proteins & receptors in brown adipose tissue (BAT) of 21-d pups born to n-3 PUFA sufficient and deficient dams. Gene expression of (**a**) *UCP1*, (**b**) *GPR120*, (**c**) FGF21 (**d**) *GLUT1*, (**e**) *GLUT4*, (**f**) *FABP3*, (**g**) *FABP4* mRNAs was measured using RT-qPCR, normalized with the mRNA expression of endogenous control, and expressed as fold changes over n-3 PUFA sufficient groups. Individual p-values are indexed in figures. Values are expressed as mean ± SEM (n=8). *p<0.05 vs. n-3 sufficient diet (Student’s t-test).

### 3.6 Expression of glucose transporters and fatty acid storage mediators in pup’s white adipose tissue

To examine if pup’s body fat accumulation was associated with the dysregulated expression of glucose transporters and lipid storage mediators, UCP1, ADRP, FABP3, FABP4, and GLUT1, GLUT4 were studied in WAT of 21-day-old pups. The UCP1 expression was undetected in the n-3 PUFA deficient WAT (data not shown). The ADRP expression was significantly increased by ∼1.67 folds (p=0.002, **Fig.5 a-b)**. However, the expression of FABP3 and FABP4 did not change between these groups (p>0.05). Like BAT, expression of GLUT1 was significantly decreased by ∼10.2 folds (p=0.0002) in the n-3 PUFA deficient WAT, whilst the expression of GLUT4 did not change between these two groups **(Fig.5 c-d)**.

**Fig. 5.**
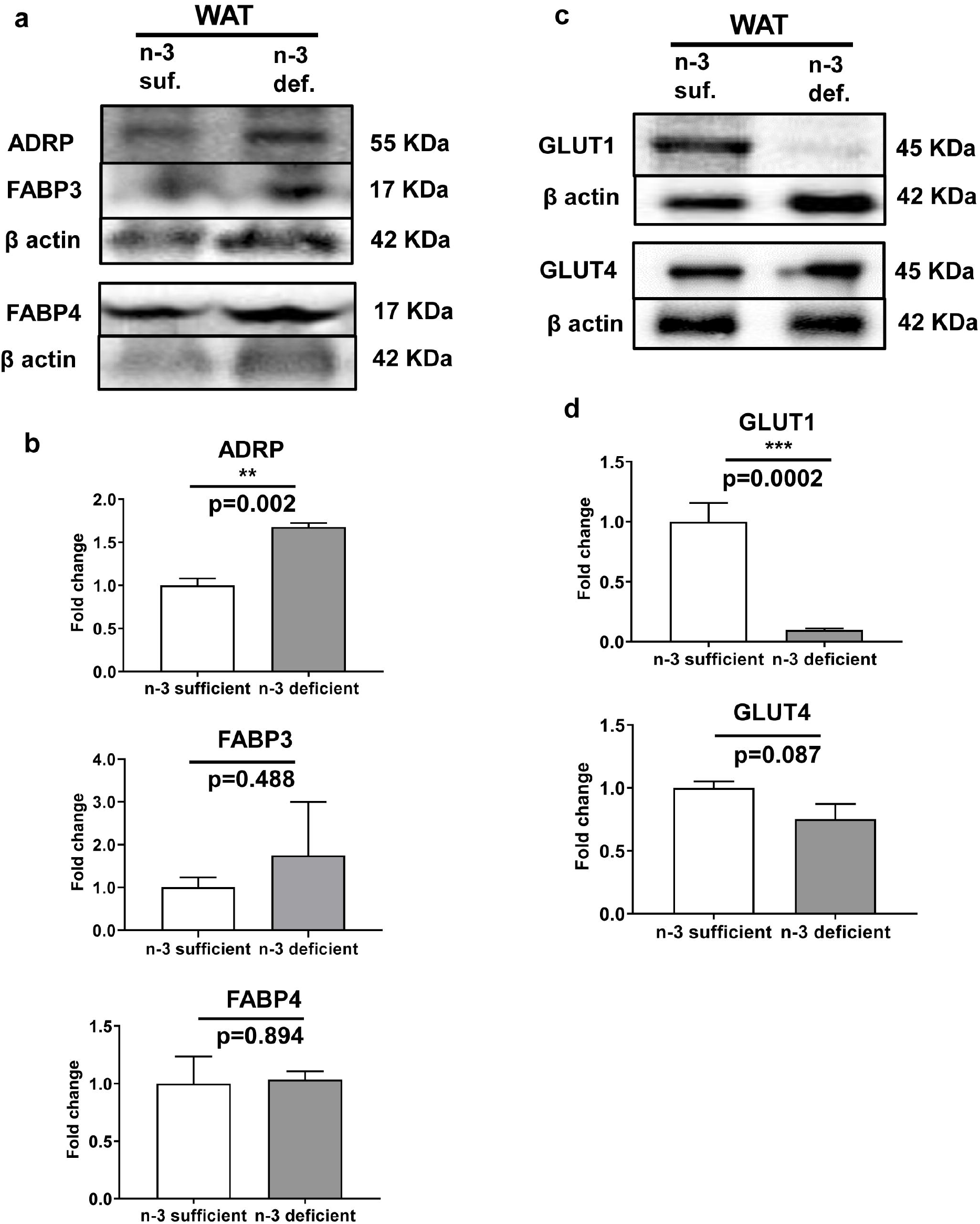
Effect of n-3 PUFA deficiency on the expression of (**a-b**) lipid storage & binding proteins and (**c-d**) glucose transporters in white adipose tissue (WAT) of 21d pups (n=4 per group). Expression levels were evaluated by immunoblotting and expressed as indicated before. Data are expressed in mean ± SEM (n=12). **p<0.005; ***p<0.0001 vs. n-3 sufficient diet (Student’s t-test).

The mRNA expression of lipogenic genes and adipose glucose uptake modulators were measured in WAT. The lipoprotein lipase (*LPL*) was upregulated by ∼1.7 folds (p=0.007), and a thermogenesis mediator *PRDM16* was upregulated by ∼1.6 folds (p=0.038), but *PGC1*_α_ expression was not affected in the n-3 PUFA deficient WAT (p>0.05). *UCP1* mRNA expression was undetected (Ct>38) in WAT. The expression of *GLUT* mRNAs (*GLUT1, GLUT3*, and *GLUT4*) was comparable (p>0.05) between n-3 PUFA sufficient and deficient WAT. The changes in the expression of WAT *IL6, PPAR*_γ_, *GPR40, GPR120*, and *ADRP* mRNAs remained insignificant between these WAT groups (p>0.05).

### 3.7 Effects of n-3 PUFA deficiency on gene expression in the liver

Since the n-3 PUFA deficiency led to an imbalance in the plasma n-6 and n-3 fatty acids, mRNA expression of desaturases, elongases, and fatty acid transporters were investigated in the liver. Expression of *FASN* (∼1.34 folds), *SCD1* (∼1.74 folds), and *MFSD2A* (∼2.0 folds) mRNAs were significantly increased (p<0.05), and *FABP3* expression was significantly decreased in n-3 PUFA deficient fed liver (**Fig.6 a-e)**. However, changes in *FABP4, PLIN2, PLIN3, FADS1, FADS2, ELOVL2, ELOVL5*, and *ELOVL6* mRNA expression were insignificant between these groups (p>0.05). Thus, an imbalance of n-6 and n-3 fatty acids in the plasma affected the expression of fatty acid transporter and metabolic genes in the liver as a result of n-3 PUFA deficient diet.

**Fig. 6.**
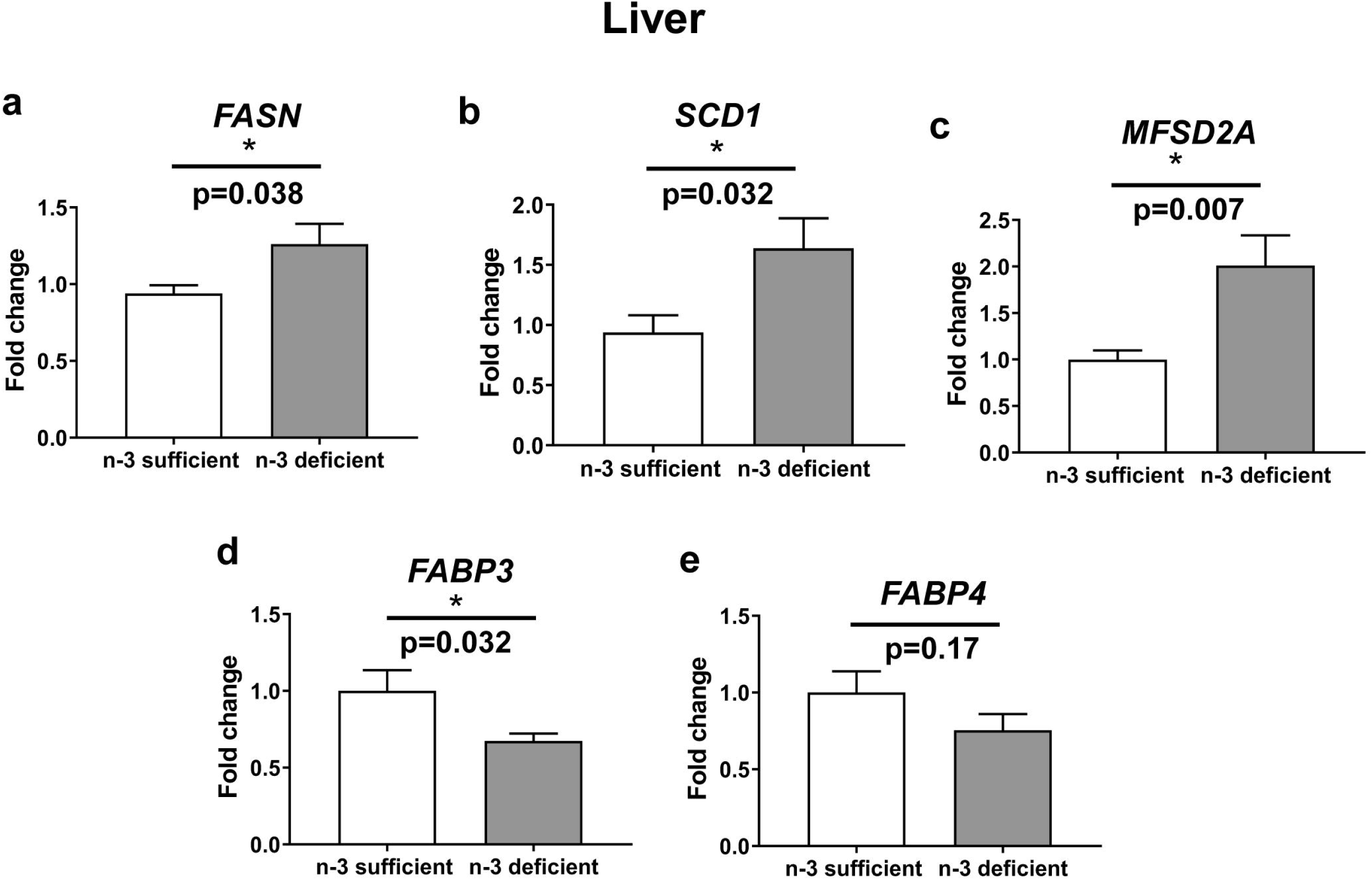
Effects of n-3 fatty acid deficiency on the mRNA expression of fatty acid metabolic and transporter genes in the liver. The mRNA expression (**a**) *FASN*, (**b**) *SCD1*, (**c**) *MFSD2A*, (**d**) *FABP3*, (**e**) *FABP4* was measured using RT-qPCR, normalized with the expression of endogenous control, and calculated as fold changes over n-3 PUFA sufficient groups. Values are expressed as mean ± SEM (n=8). *p<0.05 vs. n-3 sufficient diet (Student’s t-test).

**Fig. 7.**
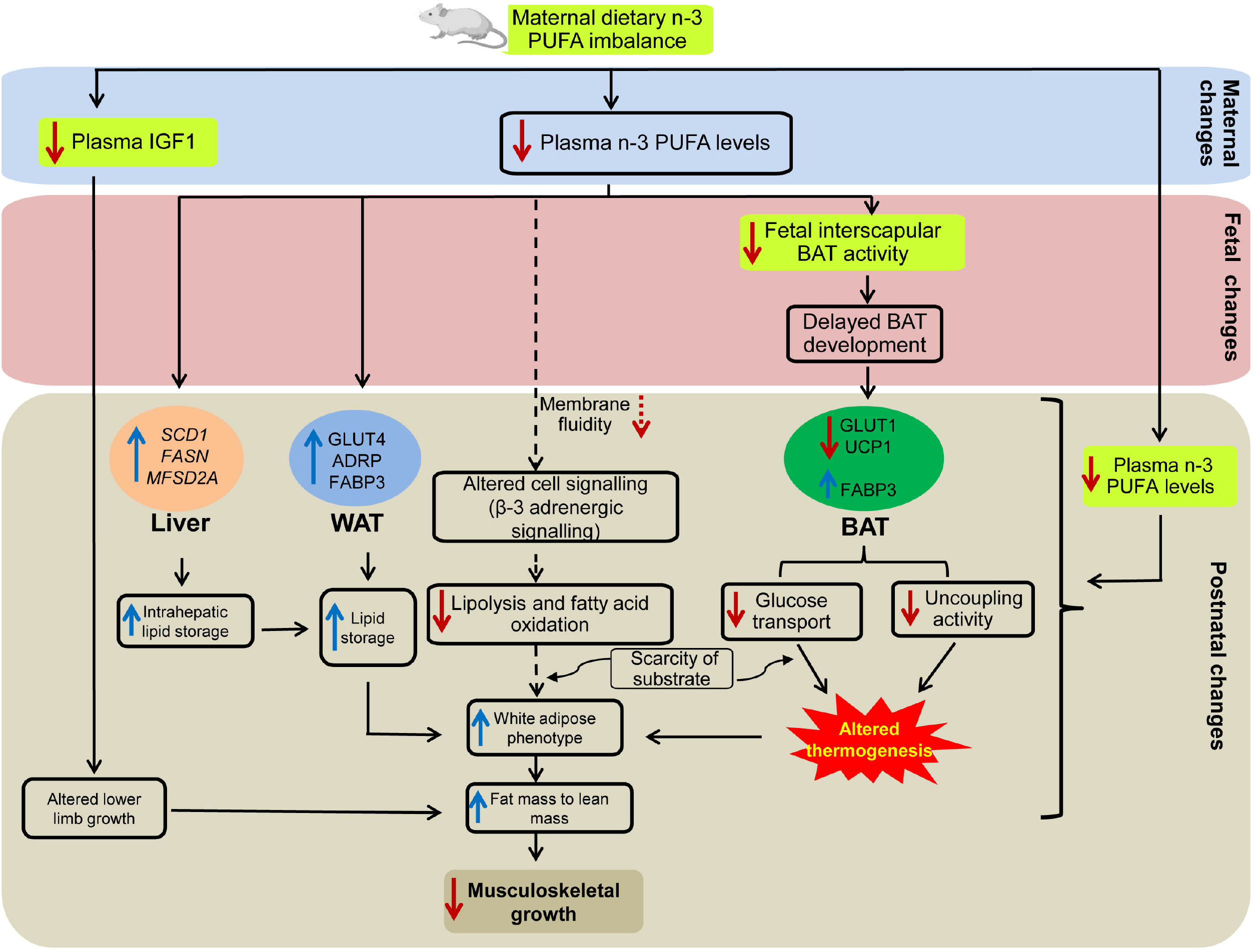
The maternal dietary n-3 PUFA imbalance and its impact on mediators associated with thermogenesis, growth hormone and metabolism at maternal, fetal and offspring levels. Based on available evidence and present data, a putative role of maternal n-3 PUFA during pregnancy and lactation in regulating fetal adiposity and offspring’s skeletal growth is postulated. Maternal n-3 PUFA modulates thermogenic development of the fetal adipose by changing the expression of UCP1, its own IGF1 levels and the expression of lipid metabolic mediators and glucose transporters in the offspring’s liver, brown & white adipose. The n-3 PUFA induces adipose thermogenesis and promotes the metabolic partition of fatty acids towards oxidation in modulating musculoskeletal growth in the mice offspring. Solid arrows indicate present data, while broken arrows define established pathways. Blue arrows indicate upregulation, and red arrows refer to downregulation.

## 4. Discussion

The study, for the first time, showed that maternal n-3 PUFA deficiency dysregulated the development of fetal adipose browning, impaired body-fat distribution and growth hormone patterns in the offspring. The offspring at the fetal developmental stages showed a reduced UCP1 expression in thermogenesis sensitive iBAT, indicating impaired BAT development might lead to increased adiposity in the n-3 PUFA deficient offspring. In addition, maternal n-3 PUFA deficiency reduced femur and tibia elongation, increased ADRP, FABP3, GLUT4 expression in BAT, increased IGF-1 levels, decreased GLUT1, UCP1, and *GPR120* expression in the offspring BAT. Despite isocaloric diets used in this study, n-3 PUFA deficiency reduced expression of UCP-1, resulting in lowered thermogenesis, which could lead to fatty acids’ metabolic mobilization towards triglyceride storage. Imbalance in the n-6 to n-3 PUFA ratio in the diet and lower quantity of n-3 LCPUFAs in the n-3 PUFA deficient mice thus might promote an obese phenotype, since n-6 and n-3 fatty acids have opposing effects on adipogenesis and energy homeostasis. The increased fat mass and stored body fatIGF-1 levels, decreased expression of BAT thermogenesis mediators, altered expression of glucose transporters, and lipid metabolic mediators collectively contribute to the offspring’s skewed musculoskeletal growth and development due to altered thermogenesis as a consequence of maternal n-3 PUFA deficiency.

In the present study, body fat accumulated in the n-3 PUFA deficient offspring is probably the result of inefficient thermogenesis. GPR120, a receptor or sensor of n-3 PUFA, is highly expressed in BAT [40] and stimulates brown fat activation by inducing FGF21 [31]. Lower induction of BAT activity and WAT browning could be due to decreased *GPR120* and *FGF21* expression observed in this study. EPA stimulates BAT thermogenesis in mice through GPR120-dependent epigenetic upregulation [9]. The absence of n-3 LCPUFAs, particularly EPA in n-3 PUFA deficient plasma (**Table 1**), observed in this study, indicate lower fatty acid oxidation and BAT thermogenesis in these mice. The n-3 PUFA deficiency-induced fat accumulation could be due to destabilized somatomedin growth homeostasis through increased IGF-1 transcription and secretion with the simultaneous activation of the JAK-STAT anabolic pathway [41]. Again, FGF21, which GPR120 activates, elicits similar growth signalling effects as circulating IGF-1 [42]. Thus, maternal n-3 PUFA deficiency can induce body fat accumulation in the offspring by regulating BAT thermogenesis involving the IGF1-FGF21-GPR120 signalling axis.

Energy homeostasis is maintained either by storage or by expenditure. Failure in these processes leads to the unintended accumulation of fats, leading to adiposity. Non-shivering thermogenesis, one of the thermogenic processes, involves UCP1 loss/decreased expression, leading to reduced thermogenesis, and increased adiposity [43]. UCP1 is a crucial regulator for adipogenesis during browning [26]. Localization and decreased expression of UCP1 in the fetus’s (age gD17) iBAT (**Fig.2**) indicates dysregulated differentiation during fetal BAT development of n-3 PUFA deficiency. Studies on mice fetal tissue revealed that iBAT is differentiated during gD17 [44]. The lowered presence of UCP1 and its reduced expression reflects the inadequate brown adipogenesis induced by thermogenic fat development, which could have increased fat mass subsequently in the offspring.

The n-3 PUFA may play a signalling role in thermogenesis by increasing BAT mass and fatty acid oxidation. A comparable n-6/n-3 PUFA (5:1) diet used previously showed a significant increase in the BAT mass might reflect a similar outcome with the n-3 sufficient (n-6/n-3 PUFA = 2:1) mice of the present study [20]. In the presence of dietary n-3 PUFA, adipocytes might acquire BAT-like phenotype within WAT and improved energy dissipation by enhancing fatty acid oxidation within this depot due to increased UCP1 expression in the fetus (**Fig 3a**) observed in this study. Fatty acid oxidation with concomitant intracellular lipolysis is pivotal for proton motive force generation that drives proton leakage via UCP1 [45]. The β-oxidation and intracellular lipolysis are stimulated by activating the β-adrenergic receptors (β3AR). Transmission of adrenergic signals is critical in the recruitment of BAT [46]. Interaction of β3AR and its downstream adenylyl cyclase is critical for the recruitment of BAT and influenced by the membrane fluidity induced by altered plasma n-6 to n-3 PUFA content [46] [47]. DHA increased β-AR expression and activated downstream signalling [48, 49] indicates its deficiency due to n-3 PUFA may negatively affect the interaction between β-AR and adenylyl cyclase signalling through decreased plasma membrane fluidity. The brown fat’s thermogenic capacity was increased by improved UCP1 and oxygen consumption when mice were fed with a PUFA-rich diet [46]. While ARA suppresses adipocyte browning, DHA counteracts the inhibitory impact of ARA by stimulating brown adipocyte development and UCP1 expression, improving the thermogenic response of BAT and inguinal WAT to β3-AR stimulation, and reducing adiposity [8, 10, 24]. In the present study, increased plasma ARA, decreased DHA level (**Table 1**), and reduced BAT-UCP1 expression (**Fig.2c** and **Fig.3a**) in n-3 PUFA deficient mice could be associated with altered plasma membrane fluidity, β-adrenergic receptors signalling, β-oxidation, and intracellular lipolysis. All these can lead to increased fat mass in n-3 deficient offspring, predisposing them to develop metabolic diseases in later life.

UCP1 modifies adipose glucose uptake and systemic glucose homeostasis independent of increased body mass [27]. We found that expression of GLUT1 decreased and GLUT4 increased in n-3 PUFA deficient fetal BAT (**Fig 3c**). The reduced UCP1 activity during n-3 PUFA deficiency could be due to lowered GLUT1 expression and increased insulin stimulative GLUT4 activity that might favor the conversion of BAT to WAT phenotype. In addition to energy homeostasis, BAT regulates glucose homeostasis via two different metabolic pathways. The active anabolic way in which glucose uptake is controlled by insulin and another is through thermogenesis by norepinephrine [50]. GLUT1 is highly expressed in BAT and positively associated with thermogenesis [51]. GLUT4 is also found in brown adipose tissue, and its localization is controlled by insulin in adipocytes [52]. As the principal player of thermogenesis, BAT can consume a significant amount of glucose from the bloodstream in addition to free fatty acids. This consumption is mediated by stimulation of β_3_-adrenoceptors which comprises the cAMP-mediated increase in GLUT1 transcription and the *de novo* synthesis of GLUT1, and the mTOR complex 2-stimulated translocation of freshly synthesized GLUT1 to the plasma membrane leading to increased glucose uptake [53].

Increased expression of lipid droplet-associated protein ADRP in n-3 PUFA deficiency (**Fig.3a**) indicates augmented differentiation of adipocytes into fully matured adipocytes [54]. ADRP stimulates fatty acid storage in cytosolic TG and promotes lipid accumulation [55]. FABP3, a fatty acid-binding protein highly expressed in heart muscle, is also associated with BAT fatty acid metabolism [56]. The expression of FABP3 was positively correlated with UCP1 expression in BAT [56], indicating a primary role of FABP3 as a fatty acid supplier in UCP1 mediated thermogenesis. Increased expression of FABP3 mRNA and protein followed by free fatty acid uptake, BAT hypertrophy in UCP1-KO mice [56-58] indicates the coordinated expression of FABP3 and UCP1 is required for adaptive thermogenesis failing of which may result in accumulation of fatty acids as triglycerides leading to differentiation of BAT eventually into WAT phenotype. In our study, FABP3 expression was increased while UCP1 expression was reduced in BAT (**Fig.3a**), probably due to compensatory response resulting from n-3 PUFA deficiency. Since FABP3 is highly expressed in muscle and BAT contains cell lineage of myoblast, increased FABP3 expression in n-3 PUFA deficiency could influence mesenchymal differentiation of BAT lineage towards pre-adipocyte precursor. In essence, these findings indicate inefficient burning of fatty acids during thermogenesis leading to skewed accumulation of lipids in BAT and altering the adiposity.

IGF-1 level indicates growth dynamics, which means the pups’ growth trajectory. IGF-1 concentrations usually increase 20^th^ week onward during pregnancy. However, n-3 PUFA deficiency significantly lowered the IGF-1 levels in dams during pregnancy (**Fig.1e**). In contrast, IGF-1 expression and concentrations were significantly increased in n-3 PUFA deficient pups (**Fig 1d and 1e**). IGF1 signalling plays a decisive role in controlling brown fat development. Dysregulated IGF1, as observed in this study, could lead to defective thermogenesis and an increase in basal metabolic rate [59] to augment body fat accumulation due to impaired metabolism of fat and muscle tissue in the offspring [3, 60]. A significant positive correlation exists between the mother’s blood IGF-1 concentration and the ponderal index of the newborn [5]. The reduced lean-to-fat mass in the offspring observed in this study could be due to reduced IGF-1 levels in the mother in n-3 PUFA deficiency. However, gender-specific multigenerational growth hormonal effects need to be investigated.

The maternal diet with n-6 to n-3 PUFA ratio (2:1) used in the present study showed an increase in femur and tibia size (**Fig.1 c**) without changing bone mineral content (BMC) in 1-mo offspring (data not presented). Earlier, a maternal diet with n-6 to n-3 PUFA (9:1) increased in femur and BMC in the 4-mo offspring [15]. Despite the differences in the n-6 to n-3 PUFA ratio and the study duration, increased femur elongation was reported in both cases. The study showed that a higher plasma DHA is positively correlated with lower bone resorption in piglets [16]. Reduced growth of femur and tibia in this study could be related to lower plasma DHA (**Table 1**) observed in n-3 PUFA deficient mice. The long-term n-3 LCPUFA intake, in particular EPA, improves the mechanical properties of cortical bone in mice [14]. However, EPA was present with undetectable levels in the n-3 PUFA deficient plasma offspring. Thus, maternal n-3 PUFA deficiency can dysregulate offspring’s skeletal growth due to the absence of EPA and the lower presence of DHA.

The high n-6/n-3 fatty acid ratio (>20) affects adiposity in diverse ways, including a change in a systemic inflammatory response and lipid metabolic energy balance. However, the n-3 PUFA deficiency did not affect pro-inflammatory IL6 expression in this study (data not presented). The experimental data pointed out those distinct metabolites of PUFAs drive the body fat gain by promoting adipogenesis. ARA’s metabolism promotes pre-adipocyte differentiation into mature adipocytes, while n-3 fatty acids inhibit this process [61]. The depletion of n-3 PUFAs significantly increased diet-induced obesity via lowering energy expenditure. Perinatal feeding of mice to a high n-6 fatty acids rich diet (LA to ALA = 28:1) over four generations results in a steady rise in the fat mass of the offspring without an increase in food consumption [62]. Additionally, ARA-derived metabolites such as PGE2 and PGF2 inhibit the conversion of WAT to BAT, which is critical for maintaining energy balance [24].

The liver is essential metabolic control of dietary fats and carbohydrate. Fatty acid synthase (FASN) catalyzes the last step in fatty acid biosynthesis and is a primary determinant of the hepatic *de novo* lipogenesis. The present study showed a significant upregulation of pro-lipogenic genes such as *FASN*, stearoyl-CoA desaturase (*SCD1*) and fatty acid transporter *MFSD2A* expression in n-3 PUFA deficient liver (**Fig.6**). The increased n-6 PUFA promoted the expression of lipogenic genes, including FASN and stimulated intrahepatic fat accumulation [63]. The reduced endogenous n-6/n-3 PUFA ratio favoured decreased production of PGE2 with a concurrent decrease in the activities of pro-lipogenic regulators such as *SCD1* and an increase in the activities of a β-oxidation promoter [64]. SCD1 regulates hepatic TG production, while PUFA regulates SCD1 transcription [65]. The n-3 LCPUFA reduces SCD1 activity [66] and lowers TG synthesis [67]. Increased SCD1 expression in n-3 PUFA deficiency could promote fatty acid synthesis, lower fatty acid oxidation and favour fat accumulation in the tissue. The increased expression of *MFSD2A* indicates a compensatory response to deliver DHA during n-3 PUFA deficiency, as MFSD2A is responsible for brain-specific DHA delivery to the fetus [33].

## Conclusions

The study is limited to the fact that it could not determine if the maternal effects were solely responsible for the changes in the offspring, as they continued breast feeding whilst mother continued on n-3 PUFA deficient diet. Again, the study could not measure musculoskeletal development in mature offspring since the earliest changes were considered right after parturition. Nevertheless, our data shows that maternal n-3 PUFA regulates thermogenic development of the fetal adipose tissue by changing the expression of UCP1. Impaired fatty acid oxidation could contribute to this process. BAT thermogenesis can be induced to combat obesity, and the roles of n-3 PUFAs could be important. Increased fat mass, IGF-1, and decreased UCP1 played an essential role in promoting adiposity and decreased WAT to BAT conversion affecting thermoregulation in n-3 PUFA deficiency. Individuals with a longer leg could have a greater musculoskeletal mass than a shorter counterpart at a given size. Skeletal muscle is the primary target for glucose disposal, necessary for glucose homeostasis. DHA level of visceral fat is affected the most compared to other organs in the body of the n-3 PUFA deficient rats [68]. Excessive body fat in Indian newborns compared to Caucasians of similar age could be due to chronic intake of n-3 fatty acid-deficient diet for several generations that might raise the endogenous n-6 fatty acid levels much higher than usual [62, 69]. Maintaining the optimal n-6/n-3 fatty acids become challenging with high intakes of n-6 fatty acids. The n-3 PUFA deficiency affects the delivery of n-3 LCPUFAs to the fetus and decreases the efficiency of endogenous conversion of n-3 LCPUFAs from their precursors.

Present data suggest that maternal n-3 PUFA deficiency affect fetal development of thermogenesis sensitive fat tissue and growth hormone levels, adiposity, energy metabolism in the offspring. All these anomalies may carry subtle risks for developing obesity and impaired musculoskeletal health later.

## Abbreviations

PUFA: Polyunsaturated fatty acid
LA: Linoleic acid
ALA: Alpha linolenic acid
LCPUFA: Long-chain polyunsaturated fatty acid
ARA: Arachidonic acid
DHA: Docosahexaenoic acid
EPA: Eicosapentaenoic acid
n-3: PUFAD:
n-3: PUFA deficiency
WAT: White adipose tissue
BAT: Brown adipose tissue
iBAT: Interscapular brown adipose
HFD: High-fat diet
AIN 93: American institute of nutrition 93
gD: Gestational day
FAME: Methyl esters of fatty acids
DEXA: Dual-energy x-ray absorptiometry
ELISA: Enzyme linked immunosorbent assay
DAPI: 4′,6-diamidino-2-phenylindole
SDS-PAGE: Sodium dodecyl sulfate-polyacrylamide gel electrophoresis
RIPA: Radioimmunoprecipitation assay
HRP: Horseradish peroxidase
ECL: Enhanced chemiluminescence
IgG: Immunoglobulin G
PBS: Phosphate-buffered saline
gDNA: Genomic deoxyribonucleic acid
cDNA: Complementary deoxyribonucleic acid
mRNA: Messenger ribonucleic acid
RT-qPCR: Quantitative reverse transcription polymerase chain reaction
Ct: Threshold cycle
SEM: Standard error of the mean
kcal: kilocalorie
BMD: Bone mineral density
BMC: Bone mineral content
CTCF: Corrected total cell fluorescence
IGF1: insulin growth factor1
FABP: Fatty acid-binding protein
GLUT: Glucose transporter
MFSD2A: Major facilitator superfamily domain-containing protein 2A
UCP1: Uncoupling protein 1
GPR120: G-protein-coupled receptor
LD: Lipid droplet
TG: Triglyceride
ADRP: Adipose differentiation-related protein
SCD1: Stearoyl-CoA Desaturase
FASN: Fatty acid synthase
PRDM16: Protein related domain containing 16
PGC1α: Peroxisome proliferator-activated receptor-gamma coactivator 1 alpha
IL6: Interleukin 6
PPARγ: Peroxisome proliferator-activated receptor gamma
FGF21: Fibroblast growth factor 21
β3AR: Beta-3 adrenergic receptor

## Declaration of competing interest

The authors declare no conflict of interest

## Acknowledgments

The study was partly sponsored by the fund received from the Department of Biotechnology, Government of India (BT/PR6946/MED), and ICMR-National Institute of Nutrition (grant number 17-BS05). In addition, Mr. Vilasagaram received a fellowship from CSIR, Govt. of India. We are grateful to Ms. Shailaja, Ms. Usha, and Dr. Suryakala Gandi for their assistance in the tissue section & staining, DEXA, and animal feeding work, respectively.

## Contribution statement

VS conducted the animal trial, sample and data collection, performed major laboratory experiments, data analysis and wrote the draft manuscript; ARM performed immunofluorescence and western blotting; SV was involved in mRNA expression by RT-qPCR; ASM was responsible for the animal trial and laboratory experiments; SRK was involved in diet preparation and fatty acid composition; AI provided inputs on study design, and reviewed the manuscript; AKDR provided critical comments and review of the manuscript. SB conceptualized the study, designed, analysed, drafted, interpreted and finalized the manuscript.

---

**Supplementary Fig 1**. The line diagram shows the study design and data collection points. Swiss albino mice fed with n-3 PUFA deficient (n-3 def) and n-3 PUFA sufficient (n-3 suff.) diets. The deficiency was induced by feeding deficient ALA/LA content in the isocaloric diet.

## Notes

### Competing Interest Statement

The authors have declared no competing interest.

### Summary of Updates

Individual authors are updated in this version

