## Supplementary figures and images for "Maternal omega-3 fatty acid deficiency affects fetal thermogenic development and postnatal musculoskeletal growth in mice"

### Sup fig1

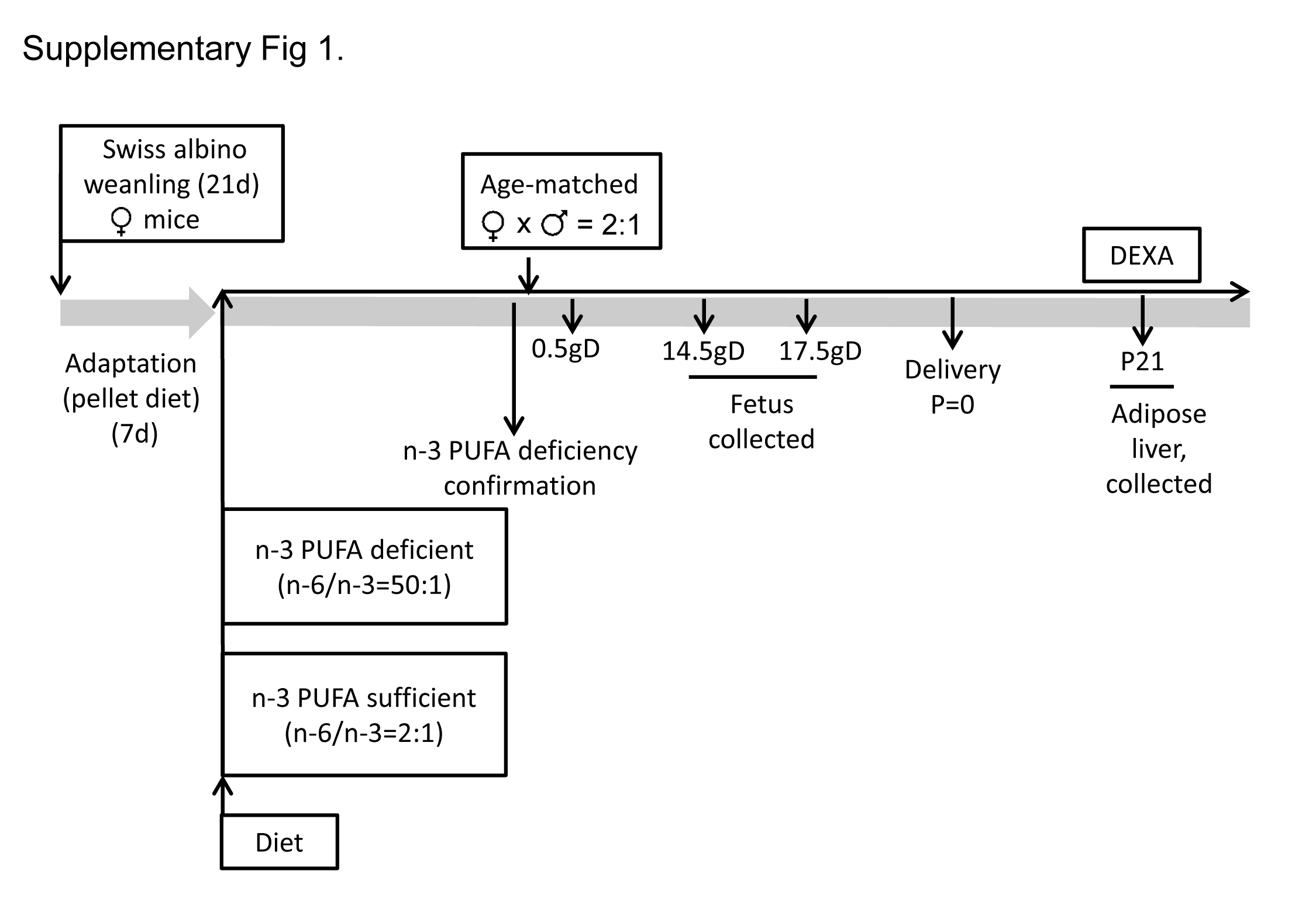
