## Supplementary material for "Maternal omega-3 fatty acid deficiency affects fetal thermogenic development and postnatal musculoskeletal growth in mice": Sup Table1

**Supplementary Table 1.** Gene pathways and predesigned SYBR green I primers used for the mRNA expression analyses

|  | Primer ID | Gene symbol | Gene ID | Gene name | Nucleotide sequences (5'-3') | Ref seq ID |
| --- | --- | --- | --- | --- | --- | --- |
| Fatty acid metabolism, storage and transporters | | | | | | |
|  | M_Fasn_1 | *FASN* | 14104 | Fatty acid synthase | F 5'-GATTCAGGGAGTGGATATTG-3'  R 5'-CATTCAGAATCGTGGCATAG-3' | NM_007988 |
|  | M_Scd1_1 | *SCD1* | 20249 | Stearoyl-CoA desaturase | F 5'-GTGGGGTAATTATTTGTGACC-3'  R 5'-TTTTTCCCAGACAGTACAAC-3' | NM_009127 |
|  | M_Fabp4_1 | *FABP4* | 11770 | Fatty acid binding protein 4 | F 5'-GTAAATGGGGATTTGGTCAC-3'  R 5'-TATGATGCTCTTCACCTTCC-3' | NM_024406 |
|  | M_Fabp3_1 | *FABP3* | 14077 | Fatty acid binding protein 3 | F 5'-AAACTCATCCTGACTCTCAC-3'  R 5'-AAAATGTCAGAGGGGAAAAC-3' | NM_010174 |
|  | M_Mfsd2a_1 | *MFSD2A* | 76574 | Major facilitator superfamily domain 2A | F 5'-ATCTACCTATTGGATGTGGC-3'  R 5'-CAGATGAGGAAGTAAGCAATG-3' | NM_029662 |
|  | M_Plin2_1 | *ADRP* | 11520 | Adipose differentiation-related protein | F 5'-ATAAGCTCTATGTCTCGTGG-3'  R 5'-GCCTGATCTTGAATGTTCTG-3' | NM_007408 |
|  | M_Plin3_1 | *TIP47* | 66905 | Perilipin 3 | F 5'-ATAGCACTAGTCTGTCATCC-3'  R 5'-TCACTGAATTTGTGATGTGG-3' | NM_025836 |
|  | M_LPL_1 | *LPL* | 16956 | Lipoprotein lipase | F 5'- GAGACTCAGAAAAAGGTCATC-3'  R 5'- GTCTTCAAAGAACTCAGATGC-3' | NM_008509 |
| Fatty acid and glucose metabolic regulators | | | | | | |
| M_Pparγ_1 *PPARγ* 19016 Peroxisome proliferator F 5'- AAAGACAACGGACAAATCAC-3 NM_001127330  activated receptor gamma R 5'-GGGATATTTTTGGCATACTCTG-3' | | | | | | |
| M_Slc2a1_1 *GLUT1* 20525 Glucose transporter 1 F 5'- AAGTCCAGGAGGATATTCAG -3 NM_011400  R 5'-CTACAGTGTGGAGATAGGAG -3' | | | | | | |
| M_Slc2a3_1 *GLUT3* 20527 Glucose transporter 3 F 5'-AGTATTCAACTCTCCACTCC-3 NM_011401  R 5-TGAAGAGGTGAAGTTAGAGG-3' | | | | | | |
| M_Slc2a4_1 *GLUT4* 20528 Glucose transporter 4 F 5'-CAATGGTTGGGAAGGAAAAG-3 NM_009204  R 5-AATGAGTATCTCATAGGAGGC-3' | | | | | | |
| Growth, receptor and inflammatory mediators | | | | | | |
| M_Igf1_1 *IGF-1* 16000 Insulin-like growth factor 1 F 5'-GACAAACAAGAAAACGAAGC-3 NM_001111274  R 5-ATTTGGTAGGTGTTTCGATG -3' | | | | | | |
| M_O3far1_1 *GPR120* 107221 Free fatty acid receptor 4 F 5'-ATCTTGATCCAAAACTTCCG -3 NM_181748  R 5-GTAAAAATGGCTCCCTTCTC -3' | | | | | | |
| M_Ffar1_1 *GPR40* 233081 Free fatty acid receptor 1 F 5'-CCATTCTGCTCTTCTTTCTG -3 NM_194057  R 5-GGGTTTATGAAACTAGCCAC -3' | | | | | | |
| M_Il6_1 *Il6* 16193 Interleukin 6 F 5'-ATTACACATGTTCTCTGGGA -3 NM_031168  R 5-TGCACTTGCAGAAAACAATC -3' | | | | | | |
| Fatty acid desaturases and elongases | | | |  |  |  |
|  | M_Fads1_1 | *FADS1* | 76267 | Fatty acid desaturase 1 | F 5'-CATACAACCATCAGCACAAG-3'  R 5'-ATGTAAGTGAAGAAGATGCG-3' | NM_146094 |
|  | M_Fads2_1 | *FADS2* | 56473 | Fatty acid desaturase 2 | F 5'-TCATCATGACAATGATCAGC-3'  R 5'-CAGGAACCTGATAAAGTTGAG-3’ | NM_019699 |
|  | M_Elovl2_1 | *ELOVL2* | 54326 | Elongation of very long chain fatty acids protein 2 | F 5'-GTTACAACTTGCAGTGTCAG-3'  R 5'-AGAAGTAGTACCACCACAAG-3' | NM_019423 |
|  | M_Elovl5_1 | *ELOVL5* | 68801 | Elongation of very long chain fatty acids protein 5 | F 5'-CTGTTCTTCCAGATTGGATAC-3'  R 5'-CCCTTTCTTGTTGTAAGTCTG-3' | NM_134255 |
|  | M_Elovl6_1 | *ELOVL6* | 170439 | Elongation of very long chain fatty acids protein 6 | F 5'-GTTTATTAATCCTCCCGCTG-3'  R 5'-CTTCTCTGGAAGTGTTTTCC-3' | NM_130450 |
| Thermogenesis mediators | | |  |  |  |  |
| M_FGF21 *FGF21* | | | 56636 | Fibroblast growth factor-21 | F 5'-CATACCCCATCCCTGACTCC-3'  R 5'-TAGAGGCTTTGACACCCAGG-3' | NM_020013 |
| M_Ucp1_1 *UCP1* | | | 22227 | Uncoupling protein 1 | F 5'-CTTTTTCAAAGGGTTTGTGG -3'  R 5'-CTTATGTGGTACAATCCACTG-3' | NM_009463 |
| M_Prdm16_1 *PRDM16* | | | 70673 | PR domain containing 16 | F 5'- ATCTACAGGGTAGAAAAGCG -3'  R 5'-TCTCCGTCATGGTTTCTATG -3' | NM_027504 |
| M_Ppargc1α _1 *PGC-1α* | | | 19017 | Peroxisome proliferative activated receptor, gamma, coactivator 1 alpha | F 5'- TCCTCTTCAAGATCCTGTTAC-3'  R 5'- CACATACAAGGGAGAATTGC-3' | NM_008904 |
| Endogenous reference | | |  |  |  |  |
|  | M_Actb_1 | *ACTB* | 11461 | Actin beta | F 5'-GATGTATGAAGGCTTTGGTC-3'  R 5'-TGTGCACTTTTATTGGTCTC-3' | NM_007393 |
|  | M_Gapdh_1 | *GAPDH* | 14433 | Glyceraldehyde-3-phosphate dehydrogenase | F 5'-AACGACCCCTTCATTGACCT-3'  R 5'-ATGTTAGTGGGGTCTCGCTC-3' | NM_008084 |
