## Supplementary material for "Maternal omega-3 fatty acid deficiency affects fetal thermogenic development and postnatal musculoskeletal growth in mice": Sup Table2

Supplementary Table 2. Body weight of foetus (14.5-17.5 gD) and 21-day old offspring

| **Weight (g)** | **n-3 sufficient** | **n-3 deficient** | **No. of mice** | **p value** |
| --- | --- | --- | --- | --- |
| Foetal weight | 3.902 ± 0.729 | 5.08 ± 0.965 | 6 | 0.3531 |
| Body weight | 6.956 ± 0.352 | 7.249 ± 0.348 | 18 | 0.5612 |
| Female pups’ weight | 6.934 ± 0.541 | 7.221 ± 0.507 | 9 | 0.7053 |
| Male pups’ weight | 6.975 ± 0.485 | 7.278 ± 0.501 | 9 | 0.6741 |

Student’s t-test between n-3 deficient and n-3 sufficient; Mean ± SEM, * p < 0.05 is considered as significant.
